# Utilizing Computational Machine Learning Tools to Understand Immunogenic Breadth in the Context of a CD8 T-Cell Mediated HIV Response

**DOI:** 10.1101/2020.08.15.250589

**Authors:** Ed McGowan, Rachel Rosenthal, Andrew Fiore-Gartland, Gladys Macharia, Sheila Balinda, Anne Kapaata, Gisele Umviligihozo, Erick Muok, Jama Dalel, Claire Streatfield, Helen Coutinho, Daniela C. Monaco, David Morrison, Ling Yue, Eric Hunter, Morten Nielsen, Jill Gilmour, Jonathan Hare

## Abstract

Predictive models are becoming more and more commonplace as tools for candidate antigen discovery to meet the challenges of enabling epitope mapping of cohorts with diverse HLA properties. Here we build on the concept of using two key parameters, diversity metric of the HLA profile of individuals within a population and consideration of sequence diversity in the context of an individual’s CD8 T-cell immune repertoire to assess the HIV proteome for defined regions of immunogenicity. Using this approach, Analysis of HLA adaptation and functional immunogenicity data enabled the identification of regions within the proteome that offer significant conservation, HLA recognition within a population, low prevalence of HLA adaptation and demonstrated immunogenicity. We believe this unique and novel approach to vaccine design that, in combination with in vitro functional assays, offers a bespoke pipeline for expedited and rational CD8 T-cell vaccine design for HIV and potentially other pathogens with the potential for both global and local coverage.

## INTRODUCTION

Since the Human Immunodeficiency Virus (HIV) was first identified, 77.3 million people have become infected of which 35.4 million people subsequently died (Sheet, Day, and People 2018). Decades of research has enabled a comprehensive understanding of the structure, genetics, mechanism of infection, immune control and immune escape to emerge, resulting in novel targets for interventions, both as therapeutic targets and for prophylaxis in the form of a broadly efficacious vaccine (reviewed (McMichael and Koff 2014)).

The structure of HIV lends itself to the development of vaccines that target the dominant surface glycoprotein gp120 and lead to the development of broadly neutralizing antibodies (reviewed by Sok and Burton (Sok and Burton 2018)). Approaches to develop immunization regimes that will bias the development of this class of antibodies to provide prophylactic protection against HIV infection are under development with the first products entering clinical assessment (Julg and Barouch 2019). However, natural control of HIV viral load following the acute viral load burst is associated with a T-cell mediated response (Altfeld et al. 2006) and this suggests that a vaccine designed to raise T-cell responses may have efficacy if it is targeted to defined antigenic regions (Ogishi and Yotsuyanagi 2019) including those with integral networked topology (Gaiha et al. 2019).

There are currently a number of T-cell vaccine candidates that utilize a variety of novel design approaches being tested in human clinical trials. The HIV Conserved vaccine (HIVCON) utilizes a conserved mosaic approach whereby regions of the proteome that have been identified as conserved within available databases are arranged in a specific regimen to both elicit T-cell responses to potential epitopes present within these regions, whilst limiting immunogenicity to the necessary joining or junctional regions (Ondondo et al. 2016). A second approach is to assemble known T-cell epitopes in a mosaic approach, whereby composite proteins are created to include common T-cells epitopes in a polyvalent design (Baden et al. 2018). A third approach, HIVACAT T-cell Immunogen, involves the construction a chimeric protein encoding 16 continuous segments of HIV derived from gag, pol, vif and nef (Guardo et al. 2016). There are pros and cons to all these approaches, but a potential caveat to utilizing conserved regions of the proteome is that historically pathogen diversity has been measured as the similarity or dissimilarity of sequences to each other, however a vaccine design should factor in how this pathogen sequence conservation is viewed by the host immune system.

Development and implementation of predictive models is becoming more commonplace as tools for candidate antigen discovery (Soria-Guerra et al. 2015). This is highly relevant for HIV vaccine discovery where there is a staggering amount of complexity posed by diversity observed within individuals (Kearney et al. 2009), within and between clades (Li et al. 2015; Taylor et al. 2008) and within populations (Maldarelli et al. 2013) making it a formidable challenge for rational T-cell vaccine design.

Here we present an *in silico* approach that complements the vaccine design strategies through the identification of HLA restricted antigenic regions within diverse HIV sequences based upon modelling of HLA restricted responses within individuals and linking these to disease progression via samples obtained from IAVI Protocol C (Amornkul et al. 2013). We show that within a population, although HLA sequences show high levels of polymorphism), there are conserved, and over represented alleles associated with the >80% of the population covered within the study. In this study, we propose the use of the artificial neural network, NetMHCpan (Nielsen et al. 2007; Nielsen and Andreatta 2016) to as a proxy to identify putative CD8 T-cell epitopes contained within the HIV transmitted founder virus (TFV) identified from the Protocol C clinical cohort of sub Saharan and East Africa. Using the transmitted founder virus sequence for relevant vaccine design is a well-established concept (reviewed here (Joseph et al. 2015)) and exploiting these predicted peptide/HLA interactions to generate additional novel metrics of HIV diversity adds another layer of information to facilitate vaccine design.

We believe that the size of the study cohort used in this investigation enables an extrapolation and scaling of the approach to global populations to enable a rationalized isolation and prediction of antigenic epitopes for any disease where a T-cell response is dominant in its control. By further informing vaccine strategies to focus the immune system against particular pathogens, incorporating potential immune recognition information into established models may increase the likelihood of success (Hare et al. submitted.).

## MATERIALS & METHODS

### Cohort characteristics

HLA profiles were evaluated from two IAVI-sponsored clinical cohorts. IAVI Protocol C is a prospective vaccine preparedness cohort studies of HIV-1 antibody negative heterosexuals or men who have sex with men in a Uganda Virus Research Institute/Medical Research Council/Wellcome Trust HIV-1 acquisition cohort study, and in a heterosexual sero-discordant couple’s cohort study in Rwanda. Subjects were given HIV counseling, condom provision and regular HIV testing either monthly or quarterly. Those who seroconverted to HIV-1 were screened for stage of primary HIV-1 infection (Amornkul et al. 2013). IAVI Protocol G was a cross-sectional cohort of ∼2000 HIV positive individuals enrolled at 13 sites around the world in order to identify circulating broadly neutralizing antibodies (Simek et al. 2009).

### Near Full Length Transmitted Founder Genomes

The selection criteria for inclusion in the generation of near full length transmitted genomes is as previously described (Baalwa et al. 2013). For this analysis, 125 Near Full length transmitted Founder genomes were evaluated from across Africa (Table 1).

**Table 1.** Distribution of input transmitted founder proteome data. Number of sequences from each country listed in parentheses

| Clade | N | Distribution |
| --- | --- | --- |
| A | 44 | Kenya (19), Rwanda (18), Uganda (6), Zambia (1) |
| C | 38 | Kenya (2), Rwanda (1), Uganda (2), Zambia (33) |
| D | 27 | Kenya (3), Uganda (24) |
| Recombinant | 16 | Kenya (6), Rwanda (4), Uganda (8) |

### HLA Distribution

The HLA binding predictor NetMHCpan was used to identify putative epitopes in 125 Transmitted Founder HIV-1 gag sequences derived from a cohort in Zambia (Claiborne et al. 2015). The distance between two sequences was defined as the percent of mismatched amino-acids in each 9mer, summed across all 9mers spanning the entire protein (i.e. a 500 a.a protein contains 492 x 9mers, each overlapping by 8 aa). This distance is dependent on sequences being aligned and therefore sequences sometimes contain gaps indicating insertions; this treats each gap character as an aa. Future analyses could consider computing an alignment-free distance.

Using this metric one can compute the distance for the entire protein or for a subset of the 9mers; the epitope-based distance included only 9mers in the alignment that were predicted to bind to at least one HLA allele. Binding was based on a threshold of 500nM, though sensitivity analyses showed similar results with different thresholds.

#### Model Implementation

For genes from each HIV virus, all 8-11mer peptides were generated. The binding affinity of each peptide to the HLA alleles described above was predicted using NetMHCpan4.1. Binding predictions were read into R and PostgreSQL for analysis. First the strain with the largest number of unique predicted binders was identified. Next, the strain that, when combined with the previously selected strains, gave the highest coverage of all predicted peptide binders was included. This strain was added to the set of selected strains and the process was repeated until all strains were included in the set. For comparison, set-building was performed a second time using randomly selected strains instead of choosing the strain that resulted in the greatest increase of peptide coverage.

#### HLA Adaptation Analysis

HLA adaptation analysis was performed as previously described (Mónaco et al. 2016). Briefly, each of the 319 peptides in the peptide set was aligned to the Zambian consensus sequence corresponding to the protein they were derived from and to HXB2. HLA adaptation was assessed using a list of statistically significant viral amino acid-HLA allele associations for Gag, Pol and Nef, previously described in Carlson et al., 2014, as well as a new list generated for Rev, Tat, Vif and Vpr based on 295 sequences derived from chronically-infected individuals from Zambia plus 237 subtype C sequences downloaded from LANL (unpublished). A peptide was adapted when either the residue was positively correlated with the HLA (referred to as adapted), or the residue was any other residue than the one negatively correlated with that HLA or the consensus (referred to as non-adapted).

#### IFN-γ ELISPOT

The predicted peptides were evaluated for ability to induce T-cell responses by IFN-γ ELISPOT using bi-specific expanded CD8 T-cells as previously described (Michelo et al). Briefly, PBMC were thawed and cultured in R10 media supplemented with IL-2 (Sigma 50U/mL final concentration) and the CD3/CD4 bispecific antibody (Genscript) to expand CD8 T-cells. On Day 7 of expansion the CD8 population was assessed by Human IFN-γ 96 well ELISPOT (Mabtech) as per manufacturer’s instructions. The peptide pools were prepared as an 11×11×11 3D matrix with each peptide occurring 3 unique pools with positive responses defined as the mean replicate count minus the mean background (mock) count where the mock controls must be <50 SFU/10^6^ PBMC and the media only wells <5 SFC/well).

#### Statistical Analysis

Data analysis was with GraphPad Prism, Python, Numpy and matplotlib. Statistical tests included Area Under Curve, Mann-Whitney test, PCoA and a Kolmogorov-Smirnov test to compare the cumulative distribution of the two data sets and computes a P value dependent on the largest discrepancy between distributions. See dataspace.iavi.org

## RESULTS

### HLA Distribution within specific Populations

HLA distribution provides an important metric describing population diversity and correlates with the breadth of viable immune recognition within that population, which is relevant to both immune protection against pathogens and vaccine design strategies. Within Protocol C, all participants were screened for HLA composition upon enrollment and Figure 1 reflects the diversity of HLA alleles within Protocol C (Amornkul et al. 2013) at a 2 field (4 digit) level of characterization (Marsh and WHO Nomenclature Committee for Factors of the HLA System 2017).This data represents the HLA diversity of 613 participants and the prevalence of the HLA A, B and C alleles is displayed as the relative percentage of the cohort.

**Figure 1.**
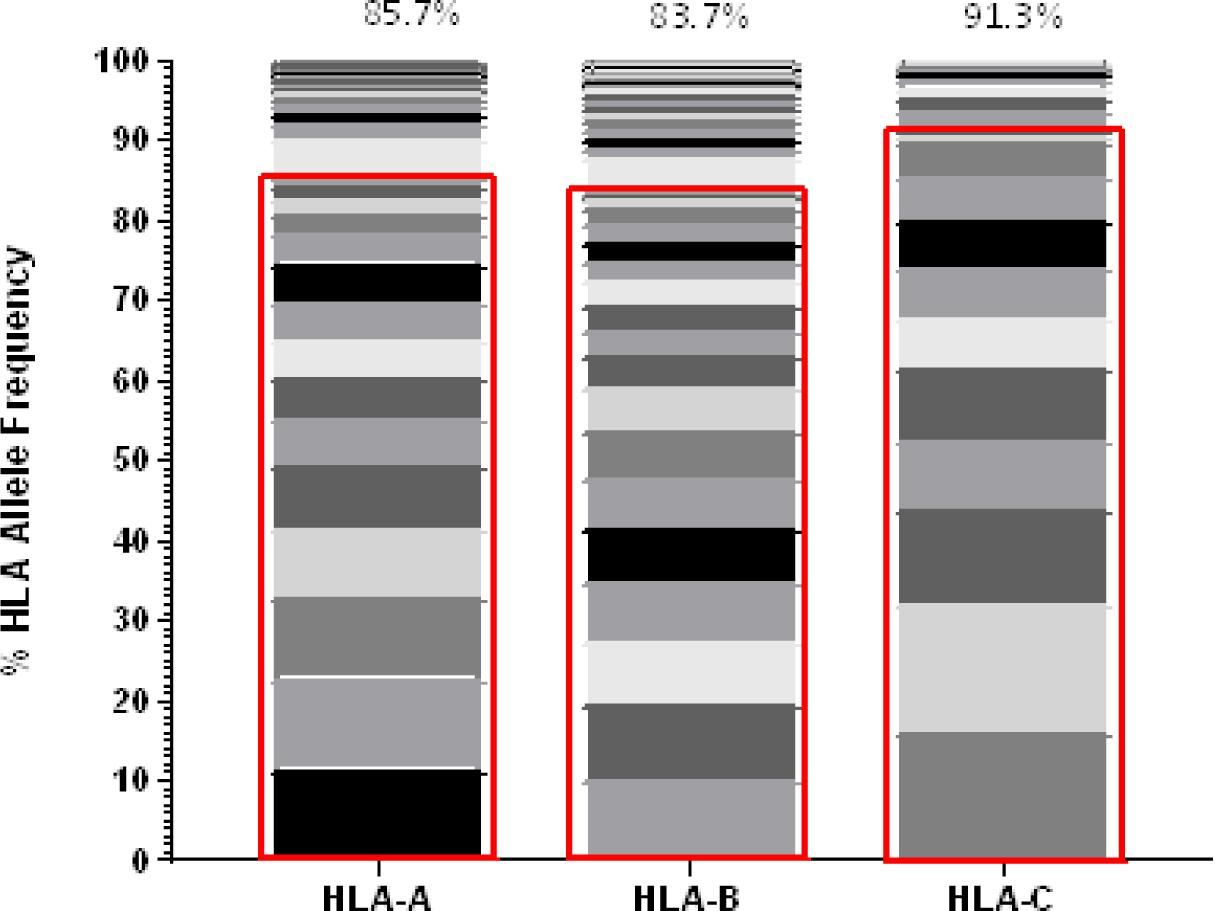
Frequency of each HLA Class I allele (HLA-A, HLA-B and HLA-C) represented within IAVI Protocol C. Alleles. Red boxes demarcate the allele frequencies contained within 13 pre-selected volunteers (Table) with percentage coverage listed above each stacked histogram plot. 17 Individual alleles contribute to HLA-A analysis, 21 Individual alleles contribute to HLA-B analysis and 13 Individual alleles contribute to HLA-C analysis

Given the expected diversity of the HLA profile, it was an unexpected observation that >80 of the HLA diversity of all alleles, are covered by 10 volunteers within the Protocol C cohort, supplemented with 3 individuals drawn from IAVI Protocol G (Simek et al. 2009) (Table 2). Furthermore, only an additional 9 alleles with frequencies >1% but <2% are excluded from this cohort (Supplementary Table 1), indicating that even with a reduced cohort size it may still be possible to capture the diversity of HLA at the sequence level.

**Table 2.** Volunteers selected for determining HLA coverage within a population

| Sample ID | HLA-A | HLA-A | HLA-B | HLA-B | HLA-C | HLA-C |
| --- | --- | --- | --- | --- | --- | --- |
| 00C175058 | A*02:05 | A*23:01 | B*07:05 | B*49:01 | C*07:01 | C*07:02 |
| 00C191996 | A*01:01 | A*03:01 | B*15:03 | B*35:01 | C*04:01 | C*06:02 |
| 00C305154 | A*68:02 | A*74:01 | B*15:03 | B*18:01 | C*02:10 | C*05:01 |
| 00C362470 | A*02:02 | A*30:02 | B*45:01 | B*53:01 | C*04:01 | C*16:01 |
| 00C305125 | A*23:01 | A*34:02 | B*08:01 | B*15:10 | C*07:01 | C*08:02 |
| 00C191735 | A*33:01 | A*74:01 | B*14:03 | B*49:01 | C*07:01 | C*08:02 |
| 00C275031 | A*23:01 | A*30:02 | B*07:02 | B*15:10 | C*03:04 | C*07:02 |
| 00C275048 | A*01:01 | A*31:04 | B*15:03 | B*51:01 | C*08:02 | C*16:01 |
| 00C365005 | A*29:02 | A*30:02 | B*42:01 | B*57:03 | C*17:01 | C*18:01 |
| 00C365007 | A*26:01 | A*29:02 | B*13:02 | B*81:01 | C*04:01 | C*06:02 |
| 00G17616 | A*02:01 | A*66:01 | B*53:01 | B*58:02 | C*04:01 | C*06:02 |
| 00G27009 | A*02:05 | A*30:02 | B*14:02 | B*58:01 | C*07:01 | C*08:02 |
| 00G27188 | A*02:05 | A*30:01 | B*07:02 | B*27:03 | C*02:02 | C*07:02 |

This analysis utilized 2-field characterization of HLA alleles, and whilst this enables frequencies of alleles to be calculated it has several limitations when considering HLA diversity/similarity. A clear limitation is that the peptide binding profile of two alleles may not be strongly associated with the similarity of their 2-field allele representation (Sidney et al. 2008). A second method for characterizing HLA allele diversity involves the assessment of the amino acid sequence of the MHC protein with a focus on the peptide binding groove (Ngumbela et al. 2008). Building on this idea, an alternative, advantageous approach to assessment of the diversity of the HLA frequency may therefore be to use computationally predicted peptide binding of the HLA alleles based on machine learning algorithms trained on functional binding data as well as the amino acid sequences of the HLA proteins (Nielsen et al. 2007).

To characterize the associated peptide:HLA diversity of the volunteers listed in Table, an HLA binding profile was modelled for each allele by predicting the binding affinity for each 9mer peptide derived from a representative panel of HIV gag amino acid sequences using the NetMHCpan4.1 binding algorithm (Nielsen and Andreatta 2016). This modelling enables us to define a binding profile of each HLA allele and each volunteer based on their HLA genotype. Based on the similarities of their binding profiles we were then able to cluster HLA alleles and/or volunteers to visualize and reassess HLA diversity. For example, a two-dimensional representation of HLA diversity in Protocol C can be generated using their pairwise HLA binding similarities and principal coordinate analysis (PCoA, Figure 2).

**Figure 2.**
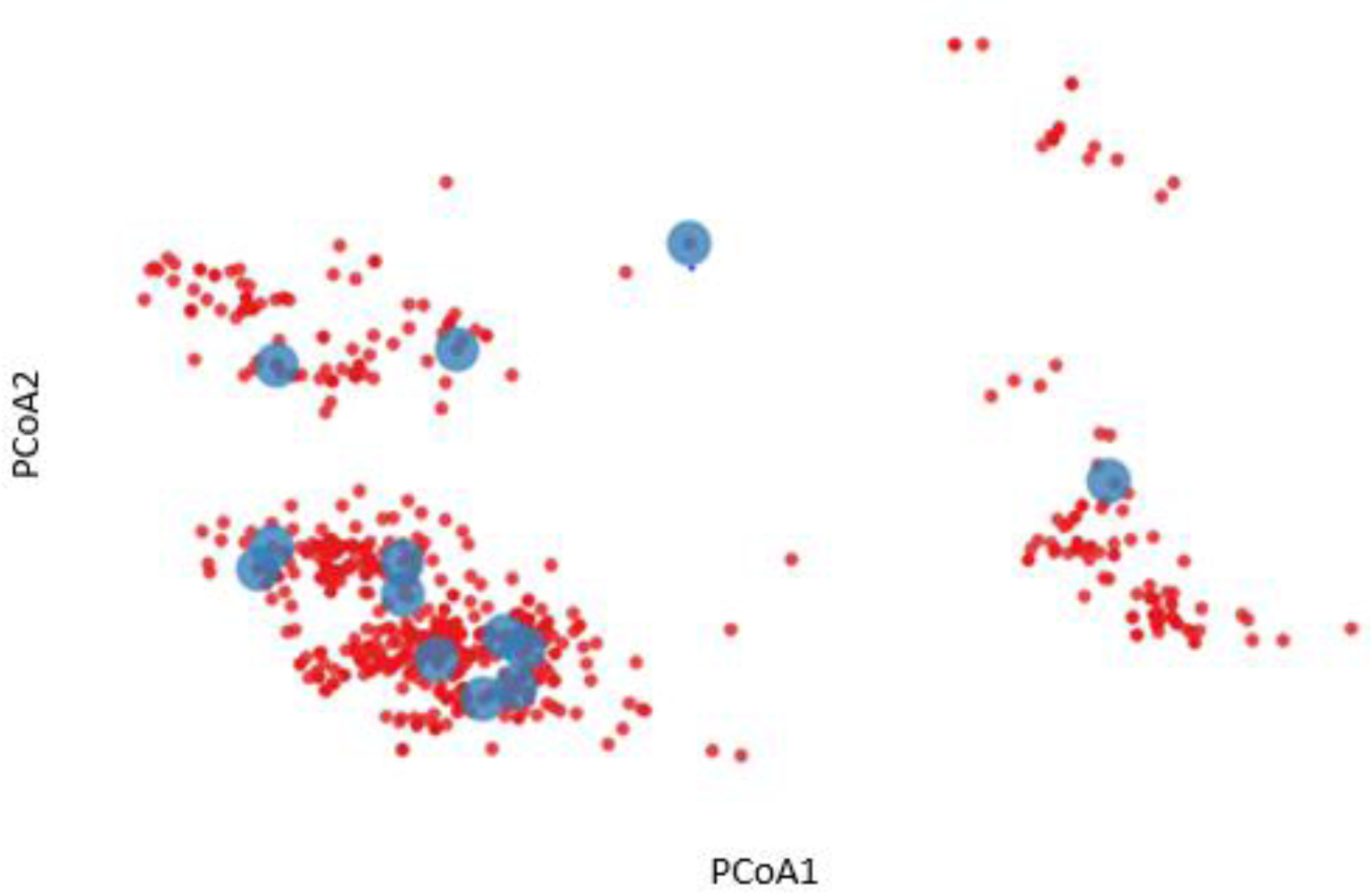
Two-dimensional representation of HLA diversity using Principal Coordinate Analysis (PCoA). A HIV-1 Gag binding profile was predicted for every HLA allele using NetMHCpan and a set of transmitted founder sequences. The binding profile of each volunteer (red dot) was defined by taking the union of predicted binding for each of their HLA alleles. PCoA was performed using the pairwise similarity matrix of all volunteers, revealing distinct clusters of individuals. A subgroup of 13 volunteers were chosen to provide optimal coverage of the HLA binding profiles (blue dots)

The analysis revealed distinct clusters of predicted HLA binding profiles which suggested that it was possible to identify a subgroup of Protocol C volunteers that were representative of the overall cohort HLA diversity.

Figure 3 illustrates that coverage of the optimal peptide sets is influenced by the prevalence of HLA alleles within the prediction. As cumulative sets of HLA alleles are removed (starting with the least frequent alleles) there is minimal loss of epitope binding coverage observed until a key inflection point is reached, leading to a precipitous loss of coverage, concordant with the frequency of the HLA alleles that are removed. Interestingly, the trend of minimal coverage loss at a minimal HLA frequency is observed independent of the size of the predicted peptide set with a comparable pattern observed for libraries of 300, 250, 200 and 150 peptides suggesting that while the HLA allele binding profile is peptide specific, it may also be independent of the peptides as long as a sufficient number are used.

**Figure 3.**
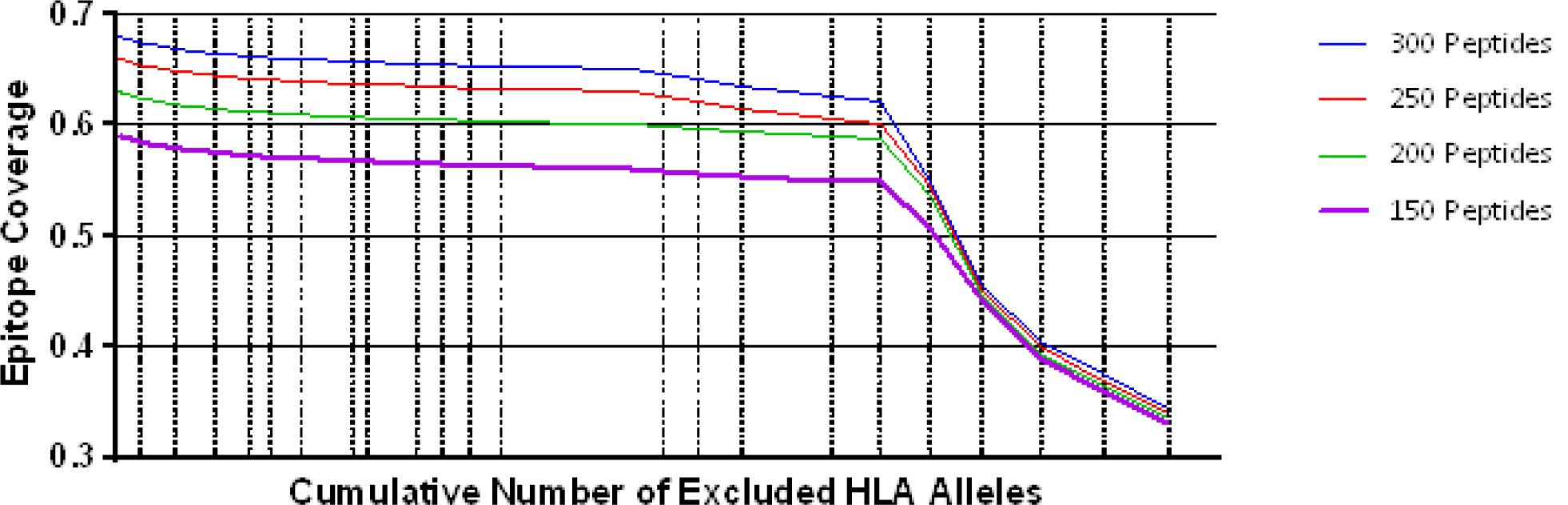
Coverage per predicted peptide calculated against a defined set of HLA alleles. Size of segments on X axis from left to right represents combined HLA allele frequencies in cohort

### Development of a predictive model for HIV diversity

Using NetMHCpan (at a 1% Binding Threshold), predicted 8, 9 and 10mer epitopes were derived from TFV gag sequences (N=*<u>127</u>*) obtained from HIV-infected volunteers enrolled in IAVI Protocol C, and identified in association with the HLA alleles present (listed in Table 1). Initial model development utilized a 1-select parameter where peptides were considered individually to determine the best coverage. This resulted in the prediction of 6562 peptides (**Error! Reference source not found**.) and no difference in best coverage mapping versus random selection (*p=0.4670*) was observed. Subsequent analysis of this model revealed that 4812 (73%) of these peptides were either unique to an individual gag sequence or present in only two gag sequences. If only peptides that were present in ≥3 virus sequences (3-select best) were considered, this led to the prediction of 1750 peptides (26.7% of the 1-select best model), which was shown to be more effective at mapping coverage than randomly selecting peptides (*p<0.0001*) (Supplementary Figure 2, Supplementary Table 2).

Further model development evaluated the effect of varying the binding threshold on the predicted outcomes. The binding threshold is a measurement of confidence that a predicted peptide will associate with the prescribed HLA, for example a 1% binding threshold factors in a 1% false positive rate. Running the model whilst varying binding thresholds at 0.5%, 1% and 2% resulted in the identification of 955, 1750 and 3023 peptides, respectively (Supplementary Table 2) No difference was observed in coverage when the 1% binding threshold was set to a less stringent 2% or a more stringent 0.5% (*p>0.9999 and p=0.6430*), therefore a 1% binding threshold was selected for all future analyses in order to maximize coverage whilst being able to distinguish additional conserved epitopes (Supplementary Figure 2)

### Modelling of HIV diversity for full length transmitted founder proteomes

These parameters were then applied to analyze 125 Transmitted Founder proteome sequences (excluding envelope) derived from IAVI’s Protocol C (see Tables 1 andTable 3 for input sample data and model parameters). The initial evaluation identified 14953 predicted peptides occurring with a frequency of 2.2% in our population. This peptide set covers all predicted affinities and coverages and may represent multiple HLA interactions/peptide. To evaluate the distribution of affinities to the primary associated HLAs with Rank Binding scores were assessed (Figure). Rank binding is an alternative metric for HLA:peptide affinity that can be deployed in order to normalize the large diversity in the range of predicted binding values for the different HLA molecules and therefore limit bias derived from over-represented HLA (Nielsen and Andreatta 2016). Rank binding assigns each peptide a score with peptides annotated as a strong binder if their score is <0.5 or a weak binder if the score is 0.5-2.0.

**Table 3.** Model Parameters

| Parameter | Values |
| --- | --- |
| Binding Threshold | 1% |
| HLA allele contributions | All HLA alleles from 13 individuals (Table ) |
| HLA haplotype weighting | 0 |
| Rank Binding | <1.0 |
| Peptide Conservation (%) | 2.2 |
| Peptide length | 8, 9, 10 & 11mers |

To further control for potential bias within the peptide-HLA interactions, the peptides were then analyzed by both affinity and Rank Binding to all predicted HLA interactions and the frequency that these peptides occurred in the population in the context of the specific HLA alleles.

This analysis identified a range of predicted binding profiles for the different peptide-HLA interactions (see Supplementary Table 1 for full HLA allele identities). HLA-A*02:02, HLA-A*31:04 and HLA-B*15:03 were identified as having particularly high predicted affinity peptide interactions, whereas HLA-B*14:03, HLA-B*15:10 and HLA-C*04:01 have much lower predicted affinity peptide interactions. This differential pattern of binding may be an artefact, explained due to the large diversity in the range of predicted binding values for the different HLA molecules. When plotted using the Rank Binding metric these differences are less pronounced although trends of stronger associations to specific HLA alleles remain.

Implementing these frequency and binding thresholds to identify HIV-specific predicted CD8 T-cell epitope peptides can be used as a functional metric to assess HIV diversity. By assuming that these predicted peptides provide a novel tool for ranking HIV proteome diversity, it is possible to assign a coverage gain value to each sequence and then utilize those values to rank each sequence for the coverage it provides within the sample population. By implementing these calculations, it is then possible to identify the sequences that are necessary to obtain the optimum level of epitope restricted sequence coverage.

The implementation of this model can then be used to target and prioritize individual proteomes. Figure 5 illustrates how for 125 transmitted founder virus proteomes, achieving 90% coverage requires 33 prioritized viruses, which decreases to 22 and 16 viruses if 80% or 70% coverage is desired, respectively (data not shown). Importantly, approximately 40% more viruses are required to achieve 90% coverage if sequences are randomly selected (n=45 p<0.0001).

**Figure 4.**
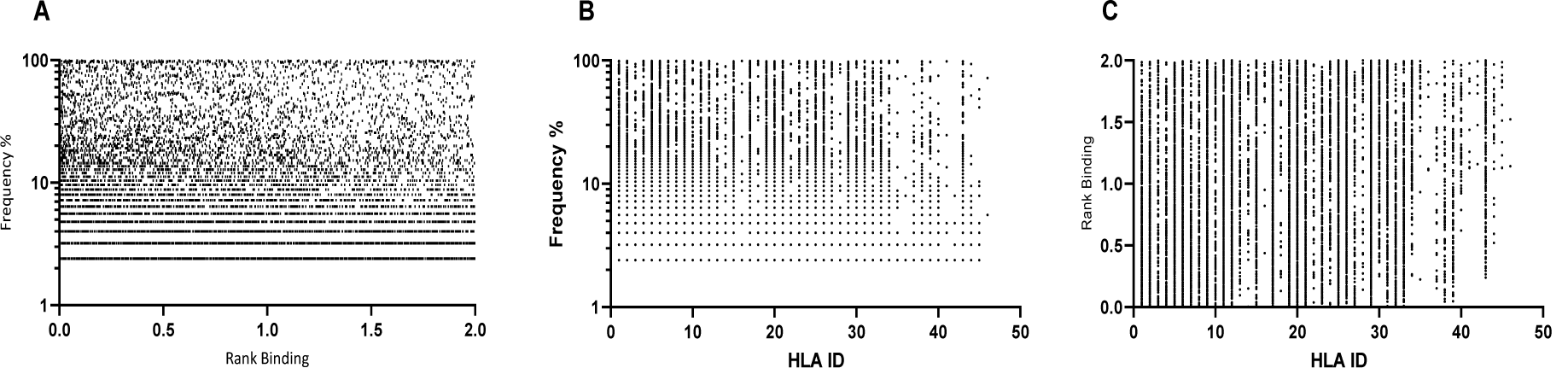
Affinity plots for all predicted peptides with conservation of ≥2.2% (n=14953). A – Predicted peptide affinity (Rank Binding) versus peptide frequency within transmitted founder proteome. B-Predicted peptide frequency versus primary associated HLA, C – Predicted peptide affinity (Rank Binding) versus primary associated HLA

**Figure 5.**
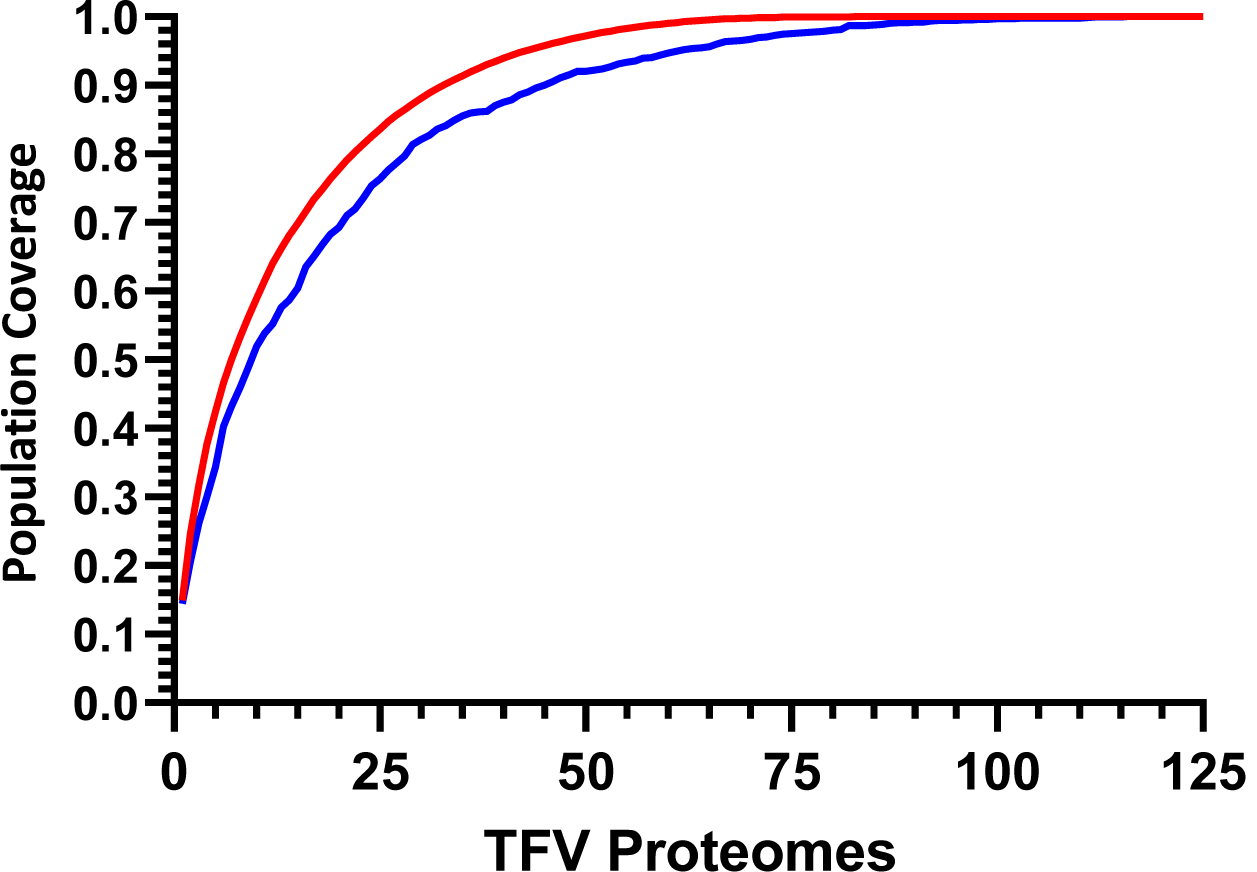
Cumulative coverage distribution plots of full length transmitted founder gag sequences using a 3-select coverage model and a 1% Binding Threshold, 3-Select best (red) and 3-Select random (blue).

### I*n silico* characterization of predicted peptides

Whilst evaluating peptides at a prevalence of ≥2.2% is desirable from the perspective of understanding population coverage, it is more challenging to map potential regions of the proteome for anti-HIV T-cell specificities due to the large levels of redundancy and overlap in evaluating each HLA/epitope interaction. By selecting HIV sequence coverage as the primary parameter and predicted affinity as a secondary characteristic the peptide library should contain both predicted high and lower affinity epitopes with optimum coverage, that may have functionality if represented at high enough abundance. Through further stratifications of the predicted peptide set to limit sequence overlap, and through assigning a minimum population coverage of 40% (selected to maintain sequence conservation and not introduce multiple sequence variations) resulted in the identification of 957 peptides. Of these peptides, an unbiased subset of 319 peptides were selected from across the proteome for further in silico and in vitro characterization.

HLA adaptation in a particular epitope is defined as the presence of a particular residue that has been statistically linked to an individual HLA, indicating a process of immune selection in that context (Mónaco et al. 2016). Vaccine design utilizing conserved epitopes may unwittingly overlook the observation that not all epitopes in the transmitted virus will be consensus and in fact, some may actively promote CTL escape (Goepfert et al. 2008). The peptides identified by the 3-select model were evaluated for predicted HLA adaptation as previously described (Mónaco et al. 2016). Of these peptides 75/332 were identified as containing a residue that was adapted, although interestingly the predicted adaptation was against alternative HLA alleles not predicted by the model for 70/75 predicted peptides with only 2 out of 5 adapted peptides associating to the primary HLA allele (Data not shown).

### Predicted peptide i*n vitro* characterization

To confirm that the selected subset of predicted peptides were recognized by anti-HIV specific T-cells, IFNγ ELISPOT assays were performed using a 3D Matrix approach described elsewhere (Fiore-Gartland et al. 2016). The peptides were evaluated in 32 HIV+ volunteers to determine the contribution of individual HLA and input sequences and correlate these metrics to observed T-cell responses.

Analysis of IFNγ ELISpot responses in HIV+ subjects who contributed their TF proteome sequence to the predicted *in silico* model revealed no significant difference in the median number of responses per volunteer (N=6) compared to volunteers that did not contribute TF sequences (median responses/volunteer N=4) (Figure 6A). Further analysis revealed that there was no bias in responses towards the volunteers with sequences predicted to contribute the most coverage versus those volunteers whose sequences contributes less to coverage (Figure 6B). Combining all the responses showed no correlation between the number of total responses/volunteer and the percentage epitope coverage offered by each peptide (Figure 6C) although the median responses/volunteer shows a trend aligning to increasing epitope coverage (data not shown).

**Figure 6.**
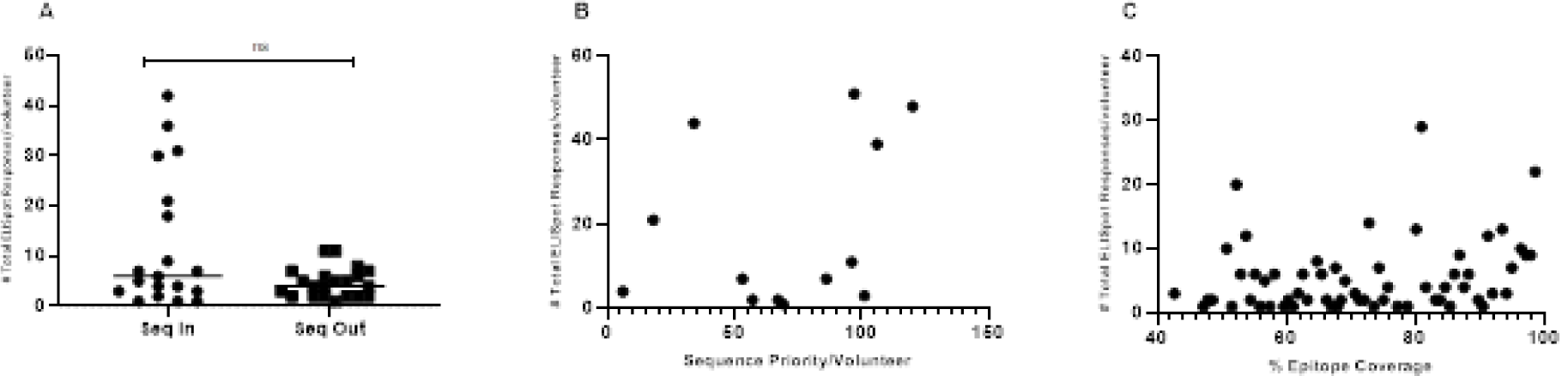
IFNγ ELISpot responses observed in HIV+ Volunteers. A – Number of total ELISpot responses observed in volunteers whose transmitted founder proteome sequence was included within the in-silico prediction (Seq In: N=19) and volunteers whose transmitted founder proteome sequence was not included within the in silico prediction (Seq Out: N=21) p=0.2104. B – Correlation of total number of ELISpot responses in volunteers whose transmitted founder proteome sequence was included within the in-silico prediction against the order of priority the sequence was predicted to occur (R^2^=0.09666 p=0.3012). C – Correlation of total number of ELISpot responses in volunteers whose transmitted founder proteome sequence was included within in silico prediction against the % coverage each epitope represented (R^2^=0.05825, p=0.0610).

## DISCUSSION

We propose that through a minimal adaptation of the existing predictive algorithm NetMHCpan, two novel parameters were defined that can be exploited to aid the rational selection of T cell vaccine immunogens. The first parameter confers the ability to assign a diversity metric to the HLA profile of individuals within a population. Although there are existing metrics for evaluating HLA profiles such as using a 2-field code or an HLA gene sequence, there are some limitations in using these parameters to assign a diversity metric score (Ngumbela et al. 2008; Sidney et al. 2008). We propose an alternative metric of HLA diversity that utilizes the predicted binding affinity of a reference amino acid sequence to assign each HLA allele an individual binding score. By evaluating the individual HLA profiles of individuals in a studied cohort, it is then possible to calculate a combined HLA diversity metric. Using these values, individual volunteers can be mapped within specific populations and distance scores calculated between each allele and each volunteer. Using this approach, we have demonstrated that it is possible to select individuals within a cohort that are “representative” of the population from which they are drawn. Implementing this stratification of volunteers may have implications for the design of smaller experimental clinical trials.

The second parameter is a metric for HIV diversity determined through the perspective of predicted binding of putative CD8 T-cell/HLA epitopes. Previous evaluations of HIV diversity rely on sequence clustering and alignments to order individual sequences. This alignment is appropriate for comparing the actual sequence of a virus genome or proteome, however this approach is limited for evaluating how an individual may recognize a specific proteome. By considering sequence diversity in the context of an individual’s HLA profile and therefore potential CD8 T-cell immune repertoire, an additional diversity metric can be layered to represent how an individual may be predicted to view a virus proteome and through combining the *in-silico* metrics, it is possible to rank HIV proteome sequences by the coverage they provide within the population across individuals. This ability to rank sequences according to putative immunogenic breadth additionally enables the interpretation of functional immunological killing assays like the viral inhibition assay (Naarding et al. 2014; Spentzou et al. 2010). Traditionally these assays have been interpreted as a binary assessment of the number of viruses inhibited. Using these novel metrics, it would now be possible to assign a population coverage score to each virus or panel of viruses and as such be able to provide an estimate as to the potential anti-virus killing activity of a volunteer based on the pattern of viruses they can inhibit.

IFNγ ELISpot analysis using the peptides predicted by the model revealed that there was no significant increase in the number of ELISpot responses/volunteer if the individual’s TFV proteome sequence was included in the prediction compared to the number of responses/volunteer if an individual’s TFV proteome was not included. This data indicates that using a subset of samples for prediction has not created any inward bias towards the input source but is representative of the population. The frequency of responses observed in this study for both groups are lower than those previously reported (Kunwar et al. 2013; Mothe et al. 2012; Sunshine et al. 2014), however this reflects the increased stringency incorporated into the development of this peptide set whereby only peptides with a predicted coverage greater than 40% were included. By way of comparison, the conservation threshold for the peptides evaluated by Kunwar *et al*. and Sunshine *et al*. were 15% and 5%, respectively, with a response rate/ volunteer of 7 and 12 epitopes, respectively (Kunwar et al. 2013; Sunshine et al. 2014).

This hypothesis indicates that through understanding the conservation, adaptation and functional score assigned to any population of target sequences, it is possible to embed this metric within algorithms to fully evaluate potential immunogenicity within the context of sequence conservation and HLA allele frequency and may contribute to expedited vaccine design and iterative testing strategies aimed at inducing protective CD8 mediated T-cell immunity. The principals underpinning this approach have applicability to other disease models and geographies for which comparative input data is available and protective CD8 responses are desirable.

## Author Contributions

EM and JH wrote the manuscript and provided conceptual input and data analysis. AFG, RR, DM, DM, LY and MN provided technical expertise and contributed to manuscript. JD, HC and CS performed ELISPOT assays EH, JG provided key supervision and support.

## Acknowledgements

This work was funded in part by IAVI and made possible by the support of the United States Agency for International Development (USAID) and other donors. The full list of IAVI donors is available at http://www.iavi.org. The contents of this manuscript are the responsibility of IAVI and do not necessarily reflect the views of USAID or the US Government.

