## Supplementary Figure 1 for "Utilizing Computational Machine Learning Tools to Understand Immunogenic Breadth in the Context of a CD8 T-Cell Mediated HIV Response"

**Supplementary Tables & Figures**

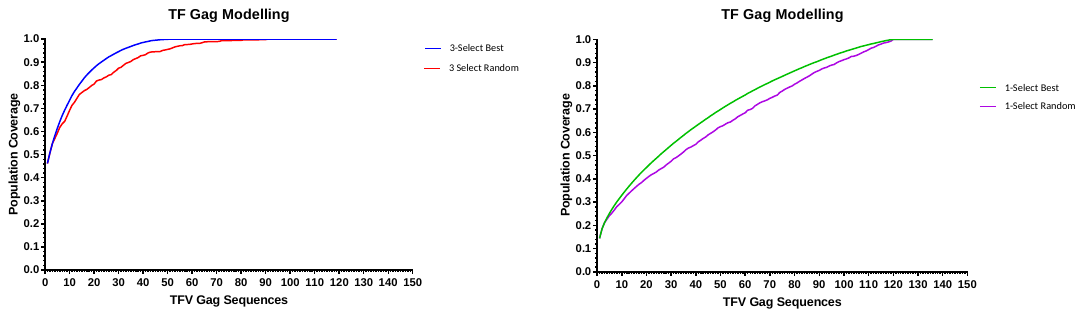

Supplementary Figure 1 Comparison of 3-select and 1-select models. Cumulative coverage distribution plots of transmitted founder gag sequences using either a 3-select coverage model (3-Select best – blue; 3-Select Random – red) or a 1-select coverage model (1-Select best – green; 1-Select Random –purple)
