## Supplementary Figure 2 for "Utilizing Computational Machine Learning Tools to Understand Immunogenic Breadth in the Context of a CD8 T-Cell Mediated HIV Response"

**Supplementary Tables & Figures**

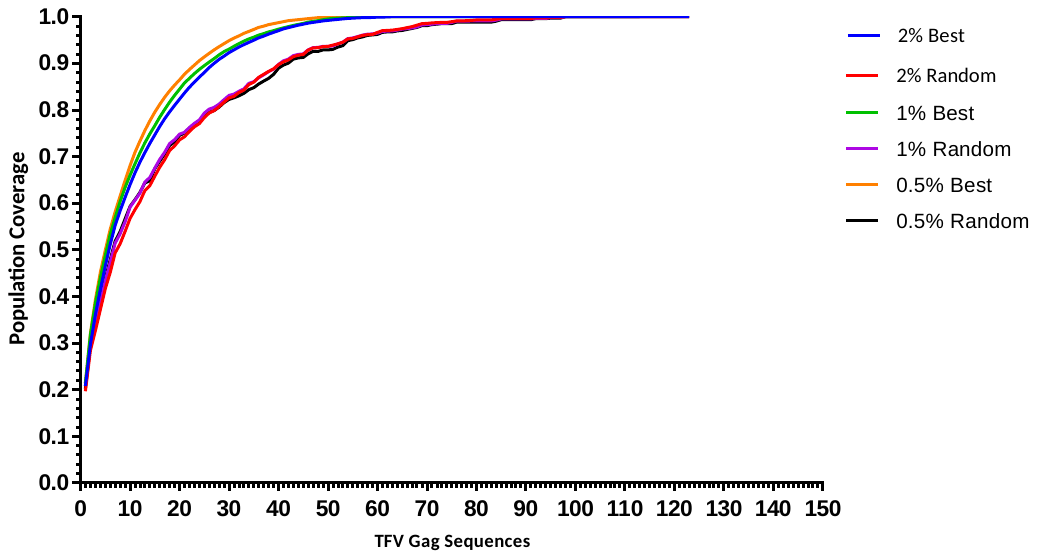

Supplementary Figure 2. Cumulative coverage distribution plots of transmitted founder gag sequences using a 3-select coverage model and varying the binding threshold. 2% Binding Threshold, 3-Select best (blue), 2% Binding Threshold, 3-Select random (red), 1% Binding Threshold, 3-Select best (green), 1% Binding Threshold, 3-Select random (purple), 0.5% Binding Threshold, 3-Select best (orange), 0.5% Binding Threshold, 3-Select random (black)
