## Supplementary Table 1 for "Utilizing Computational Machine Learning Tools to Understand Immunogenic Breadth in the Context of a CD8 T-Cell Mediated HIV Response"

| HLA-A | HLA-A Frequency | HLA-B | HLA-B Frequency | HLA-C | HLA-C Frequency |
| --- | --- | --- | --- | --- | --- |
| **A*02:01 (2)** | **11.40%** | **B*53:01 (32)** | **10.08%** | **C*06:02 (40)** | **15.98%** |
| **A*68:02 (15)** | **11.40%** | **B*15:03 (23)** | **9.34%** | **C*04:01 (39)** | **15.98%** |
| **A*30:02 (10)** | **10.17%** | **B*45:01 (29)** | **7.85%** | **C*07:01 (41)** | **11.75%** |
| **A*30:01 (9)** | **8.43%** | **B*58:02 (35)** | **7.60%** | **C*17:01 (45)** | **8.86%** |
| **A*23:01 (6)** | **8.02%** | **B*42:01 (28)** | **6.53%** | **C*02:10 (37)** | **8.86%** |
| **A*74:01 (16)** | **5.95%** | **B*07:02 (17)** | **6.20%** | **C*07:02 (42)** | **6.46%** |
| **A*29:02 (8)** | **5.04%** | **B*58:01 (34)** | **5.95%** | **C*16:01 (44)** | **6.21%** |
| **A*01:01 (1)** | **4.79%** | **B*15:10 (24)** | **5.62%** | **C*08:02 (43)** | **5.88%** |
| **A*02:02 (3)** | **4.71%** | B*44:03 | 3.97% | **C*03:04 (38)** | **5.46%** |
| **A*03:01 (5)** | **4.71%** | **B*49:01 (30)** | **3.80%** | **C*18:01 (46)** | **4.30%** |
| A*36:01 | 4.63% | **B*57:03 (33)** | **3.39%** | C*07:04 | 2.24% |
| **A*34:02 (13)** | **3.64%** | **B*18:01 (25)** | **3.14%** | C*03:02 | 1.74% |
| **A*66:01 (14)** | **2.64%** | **B*14:02 (21)** | **2.89%** | C*04:07 | 1.32% |
| **A*02:05 (4)** | **1.90%** | **B*35:01 (27)** | **2.56%** | **C*02:02** | **0.91%** |
| A*68:01 | 1.82% | **B*81:01 (36)** | **2.31%** | C*14:02 | 0.91% |
| **A*33:01 (12)** | **1.40%** | **B*08:01 (19)** | **2.23%** | C*15:02 | 0.83% |
| A*30:04 | 1.24% | **B*51:01 (31)** | **1.90%** | C*12:03 | 0.66% |
| A*24:02 | 1.16% | **B*13:02 (20)** | **1.32%** | **C*05:01** | **0.66%** |
| **A*26:01 (7)** | 1.16% | B*14:01 | 1.32% | C*16:02 | 0.25% |
| A*33:03 | 0.83% | B*44:15 | 1.24% | C*01:02 | 0.17% |
| A*01:03 | 0.58% | B*57:02 | 1.24% | C*03:03 | 0.17% |
| A*32:01 | 0.58% | B*15:17 | 0.99% | C*06:03 | 0.08% |
| A*80:01 | 0.50% | B*41:01 | 0.83% | C*15:16 | 0.08% |
| A*01:02 | 0.41% | B*42:02 | 0.83% | C*08:04 | 0.08% |
| A*30:09 | 0.33% | B*39:10 | 0.83% | C*12:02 | 0.08% |
| A*31:01 | 0.33% | B*15:16 | 0.83% | C*16:04 | 0.08% |
| **A*31:04 (11)** | **0.33%** | B*40:16 | 0.50% |  |  |
| A*02:14 | 0.25% | **B*27:03 (26)** | **0.41%** |  |  |
| A*43:01 | 0.25% | **B*07:05 (18)** | **0.41%** |  |  |
| A*66:02 | 0.25% | B*37:01 | 0.41% |  |  |
| A*02:04 | 0.17% | B*73:01 | 0.41% |  |  |
| A*11:01 | 0.17% | B*15:31 | 0.33% |  |  |
| A*26:12 | 0.17% | B*47:01 | 0.25% |  |  |
| A*23:02 | 0.08% | B*15:01 | 0.25% |  |  |
| A*26:03 | 0.08% | B*41:02 | 0.25% |  |  |
| A*29:01 | 0.08% | B*35:02 | 0.25% |  |  |
| A*31:03 | 0.08% | B*18:03 | 0.17% |  |  |
| A*66:03 | 0.08% | B*47:03 | 0.17% |  |  |
| A*74:02 | 0.08% | B*40:12 | 0.17% |  |  |
| A*74:03 | 0.08% | **B*14:03 (22)** | **0.17%** |  |  |
| A*74:05 | 0.08% | B*50:01 | 0.17% |  |  |
|  |  | B*82:02 | 0.17% |  |  |
|  |  | B*27:05 | 0.08% |  |  |
|  |  | B*07:51 | 0.08% |  |  |
|  |  | B*52:01 | 0.08% |  |  |
|  |  | B*15:47 | 0.08% |  |  |
|  |  | B*15:37 | 0.08% |  |  |
|  |  | B*15:83 | 0.08% |  |  |
|  |  | B*35:25 | 0.08% |  |  |
|  |  | B*56:01 | 0.08% |  |  |
|  |  | B*57:01 | 0.08% |  |  |

Supplementary Table 1 HLA allele frequency within Protocol C. Alleles in bold are represented within 13 pre-selected volunteers. Numbers in bold italic parentheses correspond to primary associated HLA allele identification in Figure 4
