## Supplementary Table 2 for "Utilizing Computational Machine Learning Tools to Understand Immunogenic Breadth in the Context of a CD8 T-Cell Mediated HIV Response"

**Supplementary Tables & Figures**

|  | # Peptides | AUC | | X=1, y= | | P Value |
| --- | --- | --- | --- | --- | --- | --- |
|  |  | Best | Random | Best | Random |  |
| 1-select | 6562 | 101 | 95.05 | 120 | 120 | 0.4670 |
| 3-select | 1750 | 111.5 | 108.3 | 51 | 91 | <0.0001 |
| 2% Binding | 3023 | 112.7 | 108.1 | 63 | 98 | <0.0001 |
| 1% Binding | 1720 | 113.4 | 108.5 | 57 | 98 | <0.0001 |
| 0.5% Binding | 955 | 114.2 | 108.1 | 50 | 98 | <0.0001 |

Supplementary Table 2. Model development comparing peptide conservation with levels of predicted binding affinities
